# The gastrin-releasing peptide regulates stress-enhanced fear and dopamine signaling

**DOI:** 10.1101/2020.12.31.424996

**Authors:** Yoshikazu Morishita, Ileana Fuentes, John Favate, Ko Zushida, Akinori Nishi, Charles Hevi, Noriko Goldsmith, Steve Buyske, Stephanie E. Sillivan, Courtney A. Miller, Eric R. Kandel, Shusaku Uchida, Premal Shah, Gleb P. Shumyatsky

**Affiliations:** Department of Genetics, Rutgers University, Piscataway, NJ, USA; Keck Center, Rutgers University, Piscataway, NJ, USA; Department of Statistics, Rutgers University, Piscataway, NJ, USA; Department of Molecular Medicine, The Scripps Research Institute, Jupiter, Florida, USA; Howard Hughes Medical Institute, Columbia University, New York, NY, USA; SK project, Medical Innovation Center, Kyoto University Graduate School of Medicine, Kyoto, Japan

## Abstract

Fear extinction is an adaptive behavioral process critical for organism’s survival, but deficiency in extinction may lead to PTSD. While the amygdala and its neural circuits are critical for fear extinction, the molecular identity and organizational logic of cell types that lie at the core of these circuits remain unclear. Here we report that mice deficient for amygdala-enriched *gastrin-releasing peptide* gene (*Grp*^-/-^) exhibit enhanced neuronal activity in the basolateral amygdala (BLA) and stronger fear conditioning, as well as deficient extinction in stress-enhanced fear learning (SEFL). rAAV2-retro-based tracing combined with visualization of the GFP knocked in the *Grp* gene showed that BLA receives GRPergic or conditioned stimulus projections from the indirect auditory thalamus-to-auditory cortex pathway, ventral hippocampus and ventral tegmental area. Transcription of dopamine-related genes was decreased in BLA of *Grp^-/-^* mice following SEFL extinction recall, suggesting that the GRP may mediate fear extinction regulation by dopamine.

**Impact statement:** Mice deficient for the amygdala-enriched *gastrin-releasing peptide* gene are susceptible to stress-enhanced fear, a behavioral protocol with relevance to PTSD, and show a decrease in dopamine-related gene transcription.

## Introduction

Animals and humans survive in the world by adapting their initial innate behavioral responses through learning to better navigate through the environment, which includes an exposure to threat (Kandel, Dudai, & Mayford, 2014; Tinbergen, 1951). When environmental stress and threat are excessive, the organism becomes more susceptible to trauma, generating exaggerated and unbalanced neural responses, often leading to prolonged fear (Herry et al., 2010; Maren & Holmes, 2016). Animals perceive threat as innate or learned (LeDoux, 2000, 2014). Learned (conditioned) fear is acquired after a neutral conditioned stimulus (CS) becomes associated with an aversive innate (unconditioned) stimulus (US) (Cardinal, Parkinson, Hall, & Everitt, 2002; Davis, 1997; Fanselow & LeDoux, 1999; LeDoux, 2000; Rogan, Staubli, & LeDoux, 1997). Fear conditioning in laboratory settings occurs quickly and requires only a few pairings of a CS with a shock US. Conditioned fear responses can also subdue through fear extinction. During fear extinction, because the shock US is not present, the animal learns anew that the CS no longer predicts the US, and the conditioned fear response is suppressed. Extinction does not erase the original fear memory, but creates a new memory representation.

The amygdala is a core brain region responsible for acquisition and expression of fear. However, fear extinction is dependent on the interactions of several brain regions: the amygdala, hippocampus, medial prefrontal cortex (mPFC) and some other brain regions (Luchkina & Bolshakov, 2019; Maren & Holmes, 2016). Deficits in fear extinction are also the major contributor to post-traumatic stress disorder (PTSD) in humans (Lebois, Seligowski, Wolff, Hill, & Ressler, 2019; Sangha, Diehl, Bergstrom, & Drew, 2020). Importantly, PTSD is one of the more tractable mental disorders, genetically and behaviorally, as it can be studied using rodent models of impaired fear extinction (Fenster, Lebois, Ressler, & Suh, 2018; Singewald & Holmes, 2019). There has been a significant effort to examine the neural circuits involved in fear memory, fear extinction and PTSD (Calhoon & Tye, 2015; Herry et al., 2010; Luthi & Luscher, 2014; McCullough, Morrison, & Ressler, 2016; Quirk & Mueller, 2008; Tovote, Fadok, & Luthi, 2015). However, the genetic characterization of these neural circuits remains largely unresolved (Duman & Girgenti, 2019; Lonsdorf & Kalisch, 2011; Singewald & Holmes, 2019). Defining the logic by which genes operate on specific cell types and in turn direct the neural output is therefore a central issue in understanding etiology of prolonged fear. It may also be crucial to individualized approaches to diagnosis and treatment of PTSD (Yehuda et al., 2015).

The *gastrin-releasing peptide* (*Grp*) gene may be one of the molecules critical for regulating fear extinction. The *Grp* gene is enriched in the excitatory neurons in the amygdala-associated neural circuitry of learned fear (Shumyatsky et al., 2002). When the principal neurons fire, GRP is released from excitatory neurons and binds to the GRP receptor (GRPR) expressed exclusively by GABAergic interneurons (Cao, Mercaldo, Li, Wu, & Zhuo, 2010; Kamichi et al., 2005; Lee et al., 1999; Martel et al., 2012; Shumyatsky et al., 2002). Interneurons release GABA, which via binding to the GABA-receptors decrease excitability of principal neurons. Upon binding the GRP, the GRPR stimulates interneurons to release more GABA, increasing inhibition of principal neurons and thus providing a negative feedback or feedforward loop to principal neurons. The lack of the GRPR leads to an increase in long-term potentiation in the basolateral nucleus of amygdala (BLA) as well as enhanced fear memory and deficits in fear extinction, but normal innate fear (Martel et al., 2012; Shumyatsky et al., 2002). Further demonstrating the role of the GRPergic neural circuitry in responses to threat and danger, a disruption in synaptic trafficking in GRP-positive cells leads to deficits in fear conditioning (Martel et al., 2016; Uchida et al., 2014). Moreover, administration of the GRP itself as well as GRPR agonists and antagonists affects LTP, fear memory and stress responses (Bedard, Mountney, Kent, Anisman, & Merali, 2007; Roesler, Kent, Luft, Schwartsmann, & Merali, 2014; Shumyatsky et al., 2002).

To gain insight into the role of the GRP in coping with stress and threat and to visualize its neural circuitry, we generated the GRP knockout (KO; *Grp^-/-^*) mice in which the cDNA for the enhanced green fluorescent protein (GFP) was knocked-in into the *Grp* gene locus. The *Grp^-/-^* mice were examined using a variation of Stress-Enhanced Fear Learning (SEFL) (Maren & Holmes, 2016; Rajbhandari, Gonzalez, & Fanselow, 2018; Rau, DeCola, & Fanselow, 2005), where a short acute stress is followed by fear conditioning and fear extinction, and which may be useful in understanding the mechanisms of PTSD (Sillivan et al., 2017). The *Grp^-/-^* mice demonstrated an increased susceptibility to SEFL. Moreover, we found that RNA transcription of several genes involved in the dopamine signaling and previously associated with PTSD was decreased following the recall (memory retrieval) of SEFL in the BLA of the *Grp^-/-^* mice. Retrograde viral tracing in combination with *Grp*-GFP labeling in the *Grp^-/-^* mice showed that the GRP-positive cells in the BLA receive dopamine projections from the ventral tegmental area (VTA), which is the main source of the dopamine synthesis and, as we show, contains neurons positive for both GRP and dopamine. Dopamine, a critical neurotransmitter well known to be involved in reward memories, has recently emerged as critical for fear extinction (Abraham, Neve, & Lattal, 2014; Badrinarayan et al., 2012; Gerlicher, Tuscher, & Kalisch, 2018; Hikind & Maroun, 2008; Holtzman-Assif, Laurent, & Westbrook, 2010; Kalisch, Gerlicher, & Duvarci, 2019; Luo et al., 2018; Salinas-Hernandez et al., 2018). Thus, it is possible that the GRP is critical for regulating dopamine signaling during fear extinction.

## Results

### Generation of *Grp^-/-^* mice and distribution of the *Grp*-driven GFP signal in the mouse brain

We developed *Grp^-/-^* mice in which the majority of exon 1 of the *Grp* gene was deleted and replaced with the enhanced green fluorescent protein (GFP) open reading frame (***Figure 1A; Figure 1—figure supplement 1A-C***). The *Grp^-/-^* mice of both genders develop normally and show no gross abnormalities throughout the body, including the brain (***Figure 1—figure supplement 1D***). Staining with anti-GFP antibody showed that the *Grp* gene promoter driven GFP is highly enriched in the lateral (LA) and basomedial (BMA), but not basal (BA), nuclei of the amygdala as well as in the ventral CA1 area of the hippocampus and medial prefrontal cortex (mPFC; ***Figure 1B-E***). In addition, we observed GFP expression in the retrosplenial cortex, dorsal subiculum, auditory cortex, and the entorhinal/perirhinal cortex, similar to the endogenous *Grp* expression previously examined by RNA *in situ* hybridization (Martel et al., 2012; Shumyatsky et al., 2002). Therefore, expression of the GFP knocked-in in exon 1 of the *Grp* gene is very similar to the endogenous GRP expression.

**Figure 1.**
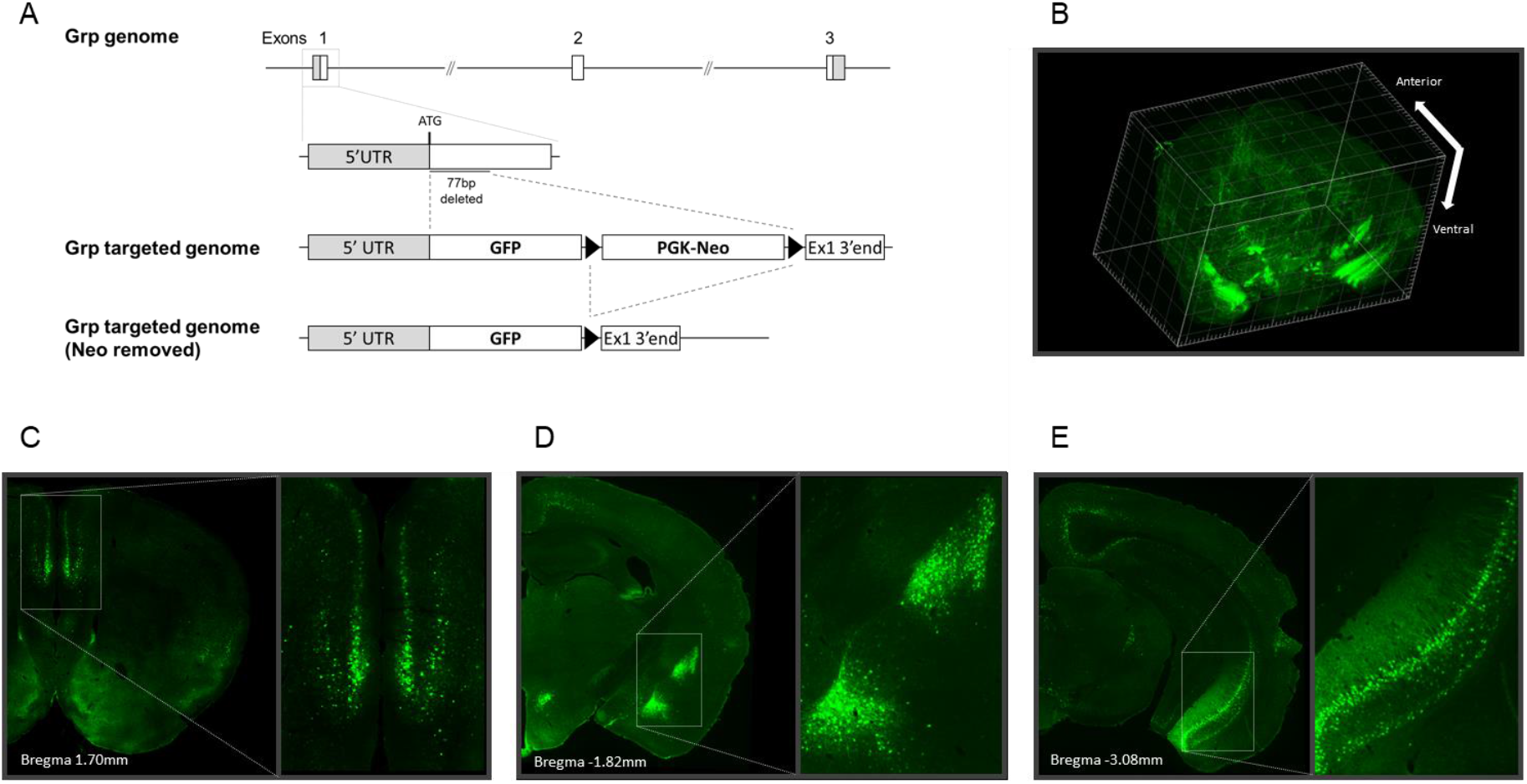
The generation of the *Grp^-/-^* mouse and the distribution of the *Grp*-gene-promoter-driven GFP signal in the mouse brain. (A) Schematic diagram of the *Grp* gene knock out. Part of exon 1 of the *Grp* gene (77bp from translation start site) was removed, and GFP ORF and neomycin cassette were inserted. The neomycin cassette was later removed by FLP-mediated excision *in vivo*. (B) The 3-dimensional map of the GFP signal in the *Grp^-/-^* mouse brain. The GFP is mainly expressed in the mPFC (C), basolateral amygdala (D) and ventral hippocampus (E).

To map the GRPergic neurons onto the amygdala-associated neural circuitry, we injected retrograde neuronal tracer rAAV2-retro-CaMKII-tdTomato (rAAV2) (Tervo et al., 2016) in several brain regions in *Grp^-/-^* mice. When rAAV2 was injected in the TE3 area, the retrograde labeling co-localized with the GRP-positive cells in the MGm/PIN area of the auditory thalamus (***Figure 2A***). When rAAV2 was injected in the lateral nucleus of the amygdala (LA), the areas labeled with tdTomato were the TE3 area of the auditory cortex (***Figure 2B***) and the MGm/PIN (***Figure 2—figure supplement 1***), two major regions sending projections to the LA (LeDoux, 2000; Pitkanen, 2000). However, only the TE3 area showed co-localization of the tdTomato and GFP (***Figure 2B***); the MGm/PIN area had no

**Figure 2.**
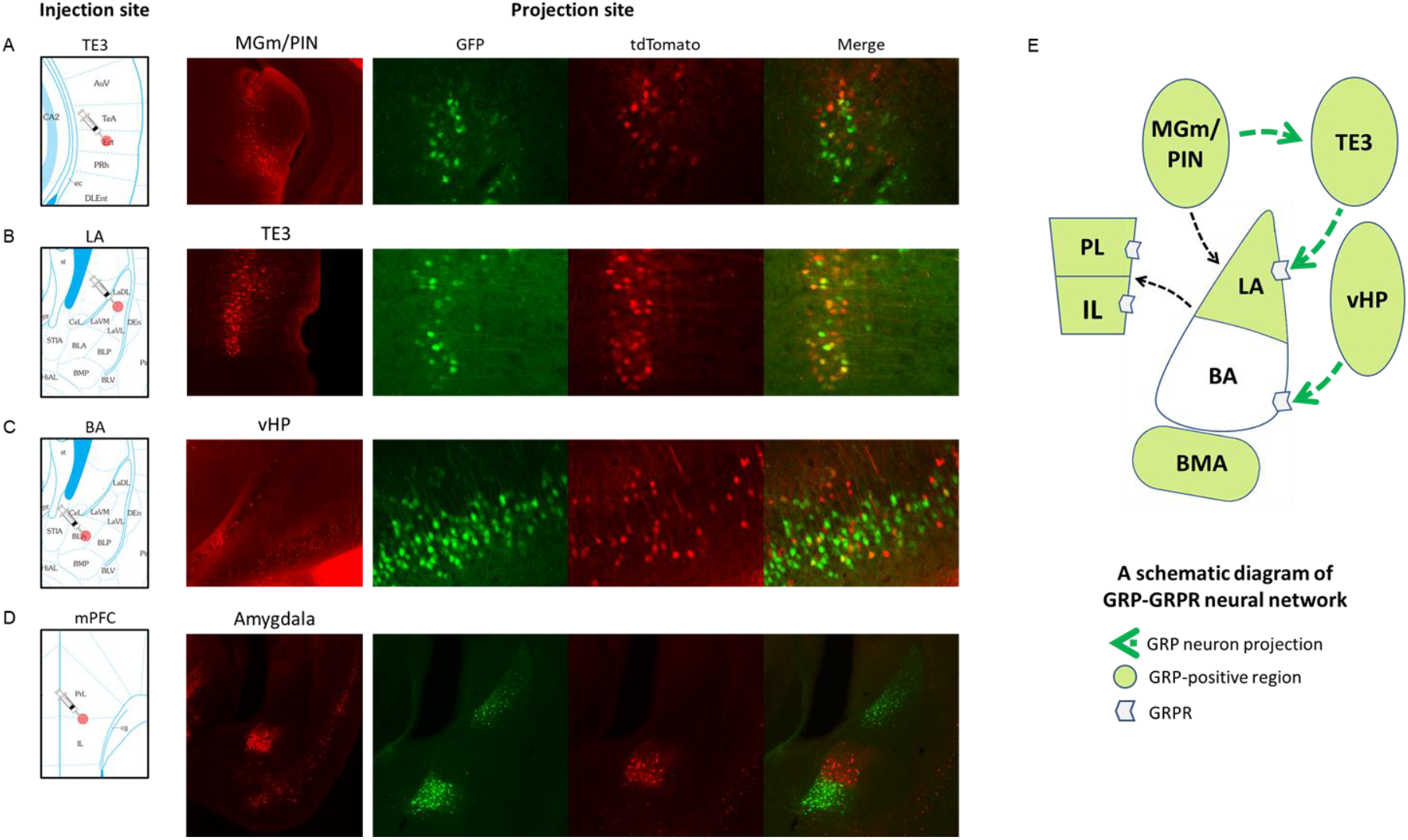
Retrograde tracing of the GRPergic projections using the *Grp^-/-^* mouse. (A-D) rAAV2-retro-CaMKII-tdTomato was injected into (A) the TE3 area of the auditory cortex, (B) lateral nucleus of the amygdala (LA), (C) basal nucleus of the amygdala (BA) or (D) mPFC of the *Grp^-/-^* mice. Three weeks following injections, the mice were perfused, and the brains were coronally sectioned at a thickness of 40 μm. The left panel shows the site of injection. Representative figures illustrate the retrograde labeling (red) of the GRPergic neurons (green) from each injection site. (E) Schematic diagram of GRPergic projections within the BLA-associated neural circuitry.

GRP-positive cells co-labeled with the tdTomato (***Figure 2—figure supplement 1***). The results of the injections into TE3 and LA show that GRP-positive cells project to the LA via the indirect MGm/PIN->TE3-LA pathway but not via the direct MGm/PIN->LA pathway.

When rAAV2 was injected in the basal nucleus of the amygdala (BA), the co-localization of tdTomato and GFP was observed in the ventral hippocampus (vHP; ***Figure 2C***), the major region projecting to the BA (Pitkanen, Pikkarainen, Nurminen, & Ylinen, 2000).

To look at the BLA projections to mPFC, we injected rAAV2 in the mPFC. Interestingly, most of the BLA projections originated from the BA area, which by itself is GRP-negative but is located exactly between the two major GRP-positive amygdala areas, the LA and the basomedial nucleus of the amygdala (BMA). There was no co-localization between GRP-positive cells and those labeled with tdTomato in the BA area or any other areas of the amygdala projecting to the mPFC (***Figure 2D***). Thus, the major area of the amygdala projecting to the mPFC, the BA, is lacking GRP-positive neurons. The BA also receives the major dopaminergic projections from the ventral tegmental area (VTA) neurons, some of which are GRP-positive (see below).

These results show that the GRPergic neural circuits are expressed in a very specific circuitry the conditioned stimulus (CS) neural pathways entering the amygdala and may play a unique role in processing of sensory information related to fear memory.

### *Grp^-/-^* mice show an enhancement in cued and contextual long-term fear memory

To characterize memory of the *Grp^-/-^* mice, we examined them in the standard protocol of fear conditioning. *Grp^-/-^* mice showed stronger long-term memory (LTM) in both cued and contextual fear conditioning (***Figure 3A-C***; post-shock freezing p=0.953, contextual LTM p=0.031, cued LTM p=0.0508). We also analyzed *Grp^-/-^* mice for short-term memory (STM) testing them 1 hour after fear conditioning training using independent groups of *Grp^-/-^* mice. There was no significant difference between wildtype and *Grp^-/-^* mice in both contextual and cued short-term memory (***Figure 3D-G***; contextual test: post-shock freezing p=0.902, contextual STM p=0.722; cued test: post-shock freezing p=0.632, cued STM p=0.115). Thus, the enhancement in memory observed in *Grp^-/-^* mice is specific to long-term, but not short-term, fear memory. These results also verified that GRP itself has an important role in fear memory processing, similar to what we found in GRPR knockout (KO) mice (Shumyatsky et al., 2002). We then bred together *Grp^-/-^* mice (GRP KO) and GRPR KO mice and assessed GRP/GRPR double KO mice in the same protocol of fear conditioning. The double KO mice showed enhanced long-term contextual and cued fear memory assessed 24 h after fear conditioning (***Figure 3H-J***; post-shock freezing p=0.963, contextual LTM p=0.018, cued LTM p=0.015). The fact that the GRP/GRPR double KO animals stronger fear memory than the *Grp^-/-^* mice suggests that the GRP and GRPR have other, perhaps secondary, ligands/receptors that they might bind, such as other members of the bombesin family (Kroog, Jensen, & Battey, 1995).

**Figure 3.**
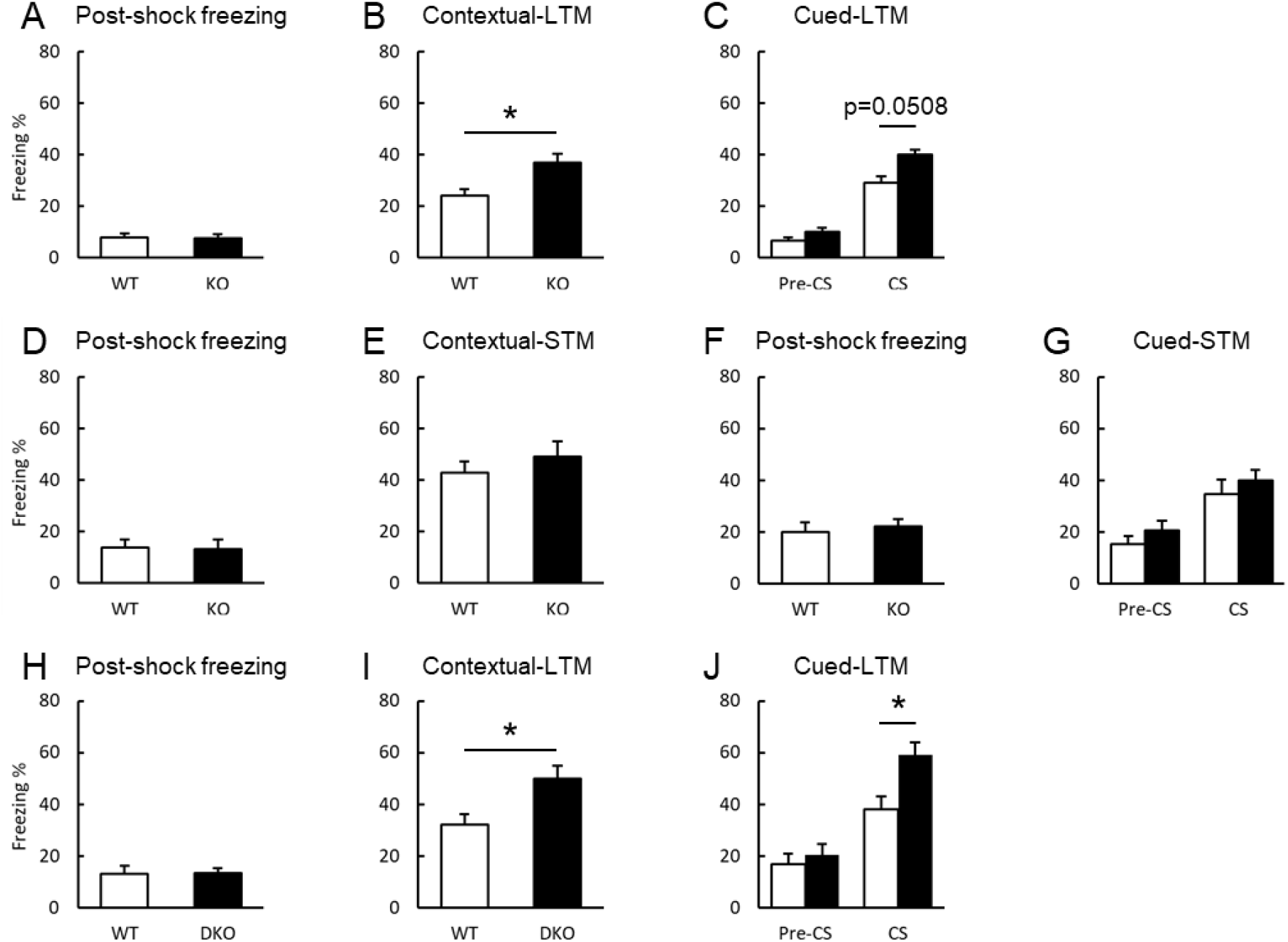
The *Grp^-/-^* mice and GRP/GRPR double knockout mice exhibit enhancement of long-term contextual and cued fear memory. A mouse was placed in a conditioning chamber for 120 sec, and a tone was applied for 30 sec that co-terminated with a foot shock (0.75 mA, 2 sec). After an additional 30 s in the chamber, the mouse was returned to its home cage. 24 hours after training (test for long-term memory, LTM), the mouse was placed back into the chamber for 180 sec (contextual test). 3 hours after the contextual test, the mouse was placed into a novel environment, and 60 sec later the tone was applied for 180 sec (cued test). Short-term memory (STM) was tested 1 hour after the training. (A-C) The *Grp^-/-^* mice showed significant enhancement of contextual and cued LTM (WT n=21, *Grp^-/-^* n=22; post-shock freezing p=0.953, contextual LTM p=0.031, cued LTM p=0.0508). (D-G) *Grp^-/-^* mice showed normal short-term contextual (WT n=10, *Grp^-/-^* n=9; post-shock freezing p=0.902, contextual STM p=0.722) and cued memory (WT n=12, *Grp^-/-^* n=13; post-shock freezing: p=0.632, cued STM p=0.115). (H-J) GRP/GRPR double knockout mice showed significant enhancement of contextual and cued LTM (WT n=10, DKO n=10; Post-shock freezing: p=0.963, Contextual LTM p=0.018, Cued LTM p=0.015). *p<0.05, unpaired Student’s t test. Data presented as mean ±SEM.

### Hyperactivity of immediate-early genes *c-Fos* and *Arc* in the amygdala of *Grp*^-/-^ mice following fear conditioning

Expression of immediate-early genes (IEG) is activity-dependent and is often used as an indicator of neural activity in the brain following memory tests or behavior. We examined RNA expression of two IEG, *c-Fos* and *Arc*, in the BLA of *Grp^-/-^* mice following single-pairing cued fear conditioning (***Figure 3—figure supplement 1***). The tissue was isolated 30 min after training in fear conditioning. Quantitative real-time PCR analysis revealed that fear conditioning-dependent expression of both *c-Fos* and *Arc* was increased in *Grp^-/-^* mice compared to their wildtype counterparts (Tukey post-hoc test, c-Fos p=0.002, Arc p=0.011). Activity of both *c-Fos* and *Arc* was normal in naïve *Grp^-/-^* mice. These results suggest that the neural activity in the amygdala of *Grp^-/-^* mice is enhanced after fear conditioning, which is consistent with their enhanced memory in fear conditioning. These results also suggest that the GRP might modulate the strength of fear memory via the signaling mechanisms that involve *de novo* gene transcription.

### *Grp^-/-^* mice show normal anxiety and pain sensitivity

We examined anxiety levels of the *Grp^-/-^* mice using the elevated plus maze (EPM), open field (OF) and light-dark (LD) box (***Figure 3—figure supplement 2A-C***; OF p=0.205, total distance p=0.785; EPM p=0.740; LD transition p=0.820). The statistical analysis showed that there was no significant difference between the genotypes in all three tests. There was a tendency for the *Grp^-/-^* mice to spend less time in the center of the open field compared to wildtype mice, but this did not reach significant difference. To verify that the increase in freezing displayed by the *Grp^-/-^* mice in fear conditioning was not due to an increased sensitivity to the shock, we examined their pain sensitivity by movement (movt), vocalization (vocal) and jump (***Figure 3—figure supplement 2D***; movt p=0.329, vocal p=0.511, jump p=0.705). There was no difference between genotypes in the intensity of the shock required to elicit these three behaviors. Thus, the increase in freezing observed in fear conditioning is due to differences in memory, but not in anxiety or pain sensitivity.

### *Grp^-/-^* mice exhibit increased susceptibility to stress-enhanced fear learning (SEFL)

To examine the *Grp^-/-^* mice in conditions where stress is combined with conditioned fear, we turned to Stress-Enhanced Fear Learning (SEFL; ***Figure 4A***) (Sillivan et al., 2017). Stress exposure consisted of two hours of immobilization/restraint, which is an acute stress (Yasmin, Saxena, McEwen, & Chattarji, 2016), followed by relatively mild tone fear conditioning (***Figure 4A-B***; two pairings of tone-shock; footshock is 0.5 mA). There was also a group of mice for each genotype, which received fear conditioning, but no shock (fear learning, FL). Then, the mice underwent fear extinction. During extinction, the *Grp^-/-^* mice froze more compared to wildtype mice (***Figure 4C***; F_1,9_=19.076, p<0.001). A repeated-measures ANOVA revealed the significant main effect of the stress on the freezing during the extinction phase in the *Grp^-/-^* mice, but not in wildtype (WT) mice (***Figure 4C**; Grp^-/-^* F_1,9_=25.860, p<0.001. WT F_1,9_=2.729, p=0.101). In the recall test performed two weeks following extinction, the effect of the genotype on freezing was observed in the SEFL group (two-way ANOVA; genotype F_1,34_=8.283, p=0.0069, stress F_1,34_=3.232, p=0.081; post-hoc: WT-SEFL vs KO-SEFL p=0.024, KO-FL vs KO-SEFL p=0.091). Following fear conditioning, the stressed group was separated into two subgroups, resilient and susceptible, based on their freezing performance during one minute of post-shock freezing (***Figure 4B***), as post-shock freezing during fear conditioning training is used as an index of stress susceptibility in this SEFL protocol (Sillivan et al., 2017). Animals that froze above the mean percent freezing in the stressed group were classified as stress-susceptible (SS), while those that fell below the mean were classified as stress-resilient (SR). Interestingly, the ratio of the susceptible group to the resilient group was higher in the *Grp^-/-^* mice compared to wildtype mice. There was a significant difference between the wildtype (WT) stress-resilient group and wildtype stress-susceptible group (***Figure 4D***; Extinction: interaction F_9,90_=2.495, p=0.013, Recall p=0.020). There was no significant difference between the knockout stress-resilient group and knockout stress-susceptible group in extinction and recall (***Figure 4E***; Extinction: interaction F_9,90_=0.455, p=0.901, Recall p=0.758). These results suggest that the increase in freezing in the *Grp^-/-^* mice results from their higher susceptibility to stress.

**Figure 4.**
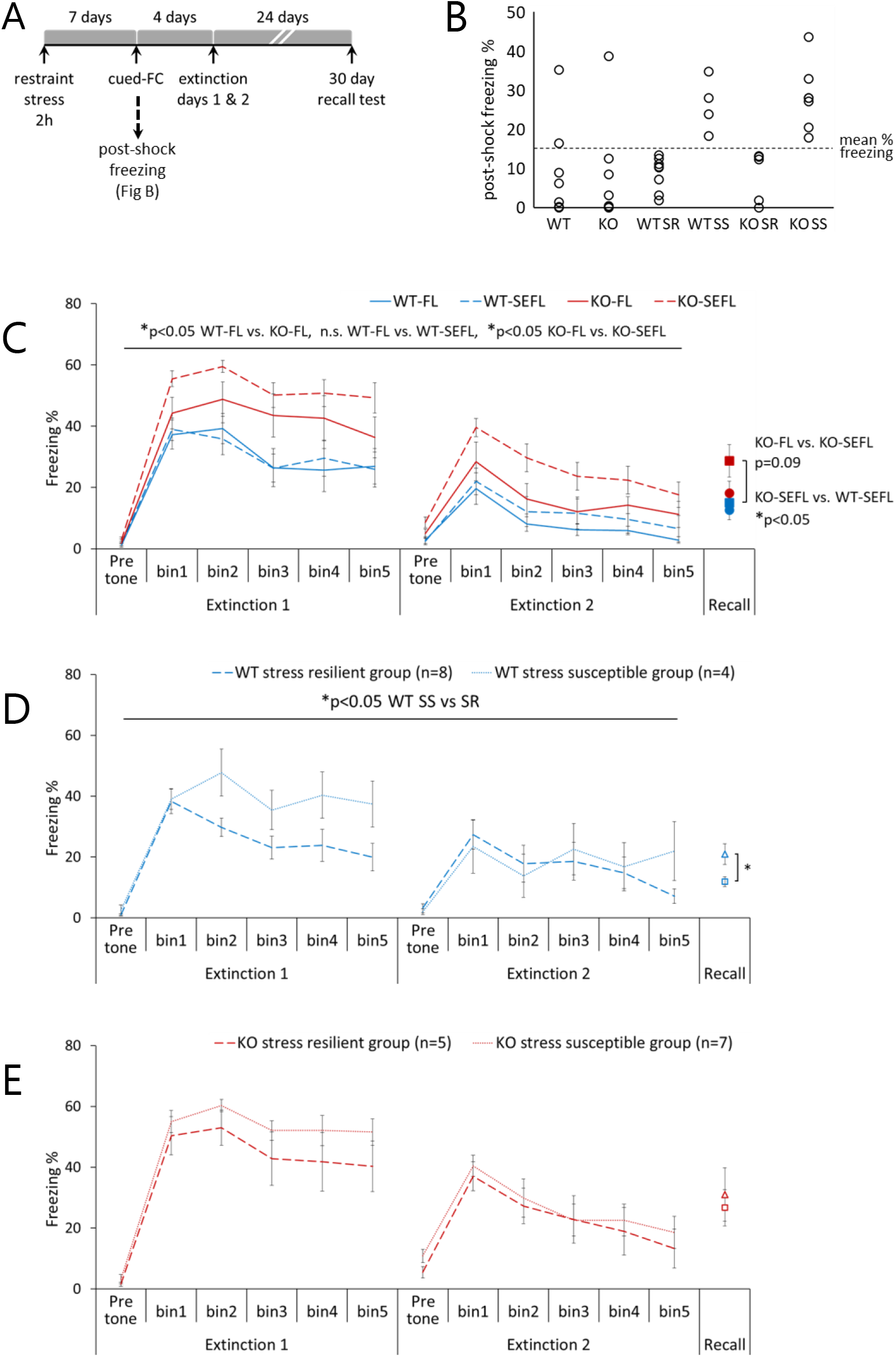
The *Grp^-/-^* mice show increased susceptibility in stress-enhanced fear learning (SEFL). (A) An overview of the SEFL paradigm. Mice underwent the following fear conditioning protocol: two minutes of exploration followed by two CS-US pairings that co-terminated with a 0.5 mA footshock (US). Extinction training (4 days post-shock) and remote memory retrieval tests (30 days post-shock) were performed in a novel context. (B) Post-shock freezing was used to separate the subjects to susceptible and resilient groups. Animals that froze above the mean % freezing for the stressed group were classified as stress-susceptible (SS), while those that fell below the mean were classified as stress-resilient (SR). (C) Course of extinction and recall test in SEFL. Shown are five bins (6 tones each) of the conditioned stimulus (CS) presentations during extinction. [WT-FL n=8,WT-SEFL n=12, KO-FL n=8, KO-SEFL n=12] (D, E) Stressed and fear-conditioned (SEFL) mice can be separated into two subgroups, stress resilient (SR) or stress susceptible (SS), based on their post-shock freezing during fear conditioning. Their extinction profiles showed significant difference between SR and SS in extinction and recall test in WT-SEFL but not in KO-SEFL. Behavioral analysis was performed using two-way repeated-measures ANOVA. Data presented as mean ±SEM.

### Induction of genes in the dopamine-signaling pathway following SEFL memory recall is dependent on the GRP

To assess the molecular events that may contribute to the behavioral phenotype of the KO-SEFL group, we used quantitative real-time PCR (qPCR) to examine transcription of several genes known to be induced following SEFL recall (Sillivan et al., 2017), PTSD and stress (Maren & Holmes, 2016; Wingo et al., 2018). We found that several genes involved in the dopamine signaling were downregulated in the *Grp^-/-^* mice compared to control mice following SEFL memory recall (***Figure 5A*** and ***Figure 5—figure supplement 2A***). These genes encode for the tyrosine hydroxylase (TH), nuclear receptor related 1 protein (NURR1) and dopamine receptor D1 (DRD1). Gene transcription of all three genes was decreased in the *Grp^-/-^* mice received SEFL, while the *Drd1* mRNA was also decreased in the *Grp^-/-^* mice that received fear extinction but no stress (FL group). There was a significant interaction between the effect of genotype and stress on a decrease in expression of the *Tyrosine hydroxylase* (*Th*) mRNA as shown by two-way ANOVA (***Figure 5A*** and ***Figure 5—table supplement 1***). Two-way ANOVA followed by post-hoc test also revealed a significant acute stress-dependent decrease of the *Nuclear receptor related 1* (*Nurr1*) mRNA expression in the *Grp^-/-^* mice and GRP knockout-dependent decrease of the *Dopamine receptor D1* (*Drd1*) mRNA expression (***Figure 5A*** and ***Figure 5—table supplement 1***). Western analysis of the TH protein showed similar levels between naive *Grp^-/-^* and control mice (***Figure 5—figure supplement 1***), suggesting that the transcription changes were induced by memory retrieval during SEFL recall. Taken together, these results suggest that the GRP may be involved in processing of stress-related memory of fear and fear extinction through the modulation of the dopamine signaling.

**Figure 5.**
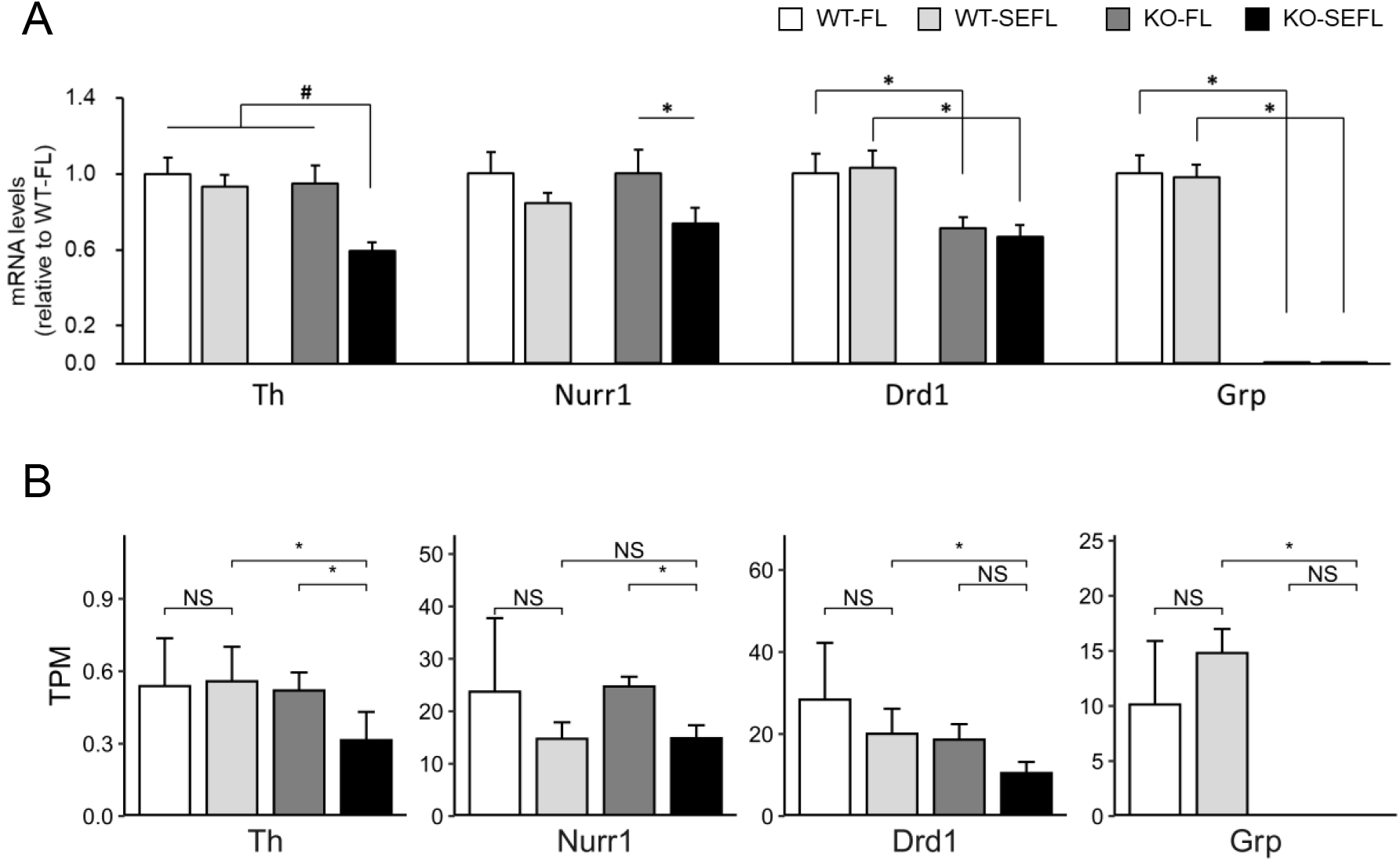
Dopamine signaling-related genes are downregulated in *Grp^-/-^* mice following recall in SEFL. Quantitative real-time PCR (qPCR) and RNA-seq analyses of dopamine signaling-related genes that were differentially expressed in the BLA. After the recall test, the mice were returned to their home cage and the amygdala tissue was dissected 30 min later. (A) qPCR analysis (WT-FL n=12, WT-SEFL n=18, KO-FL n=13, KO-SEFL n=18). All target mRNA expression levels were normalized to *Gapdh* expression and verified by normalization to *β-actin*. Results are expressed as x-fold change normalized to WT controls. All measurements were performed in triplicate. Two-way ANOVA interaction effect #p<0.05. Bonferroni test *p<0.05. Data presented as mean ±SEM. (B) Expression differences in dopamine signaling-related genes in RNA-seq data recapitulate patterns observed in qPCR datasets. Comparisons of TPM (transcripts per million) based on 4 replicates and (*) indicates significant differences based on p-values <0.05 based on a t-test.

### RNA-sequencing confirmation of the induction of the dopamine-signaling genes following SEFL memory recall

Consistent with our qPCR results, but in a separate group of animals (N=4 for each group), we found that *Th* has lower abundance (in transcripts per million (TPM)) in the KO-SEFL mice compared to all the other samples (***Figure 5B***) in our RNA-seq datasets. We also found *Nurr1* RNA abundance is lower in SEFL conditions relative to FL conditions in both the WT and KO mice, validating our qPCR results (***Figure 5B***). However, these differences are significant only for the KO strain. This is likely due to the fact that *Nurr1* abundance in WT-FL strains is highly variable. We also find that *Drd1* abundance is significantly lower in KO-SEFL relative to WT-SEFL consistent with our qPCR analyses (***Figure 5B***). Some additional dopamine-related genes showed differences in RNA-seq that we did not see in our qPCR analysis: we find that *Drd2* and *Grik2* abundances are significantly lower in KO-SEFL relative to WT-SEFL (***Figure 5—figure supplement 2B***). Interestingly, we also find abundance is significantly lower in KO-SEFL relative to WT-SEFL for the *Ppm1f* gene, which encodes for protein phosphatase, Mg^2+^/Mn^2+^ dependent 1F (PPM1F). The *Ppm1f* gene was found to be regulated by stress in mice as well as associated with anxiety and PTSD in humans (Sullivan et al., 2019; Wingo et al., 2018). While the average abundances in KO mice across both conditions are lower than in the WT mice, some of these differences are small and not significant due to high biological variability in mRNA abundances across individual samples in the WT mice. Interestingly we find that in the wildtype mice, SEFL led to no significant changes in gene-expression patterns genome-wide relative to non-stressed mice that underwent fear extinction (WT-FL; ***Figure 5— figure supplement 2C***). Similarly, under non-stress conditions (KO-FL), *Grp* gene deletion by itself failed to elicit large changes in gene-expression patterns relative to the wildtype control mice. However, when the *Grp^-/-^* mutant is treated with SEFL, we observed large and significant changes in gene-expression patterns relative to both SEFL wildtype and FL *Grp* mutant. Together this indicates that neither the stress alone nor the *Grp* gene deletion by itself has large effects but these two conditions act synergistically. Furthermore, many of the genes that were significantly affected were associated with dopamine signaling (highlighted in red), and recapitulated effects seen in our qPCR analyses.

### Overlap between cells expressing the GRP and tyrosine hydroxylase in the ventral tegmental area

To investigate the relationship between GRPergic and dopaminergic neurons, we subjected brain sections of *Grp^-/-^* mice to co-immunostaining for the tyrosine hydroxylase (TH) and GFP. TH immunostaining of the cell bodies of neurons was not observed in the BLA (***Figure 6A***), confirming previous work showing that TH is mainly expressed in neurons in the ventral tegmental area (VTA) that send projections to other brain areas including the BLA. The GFP-positive areas (LA and BMA amygdala nuclei, containing GRP-positive principal cells) within the BLA did not overlap with the BA amygdala area receiving the projections of the TH-positive neurons (***Figure 6A***). However, previous work by several groups including ours showed that interneurons positive for the GRP receptor (GRPR) are equally distributed throughout the whole BLA area (Cao et al., 2010; Kamichi et al., 2005; Lee et al., 1999; Martel et al., 2012; Shumyatsky et al., 2002). In the VTA, we found that the GRPergic GFP-positive neurons and TH-positive neurons overlap (***Figure 6B***). Combined, our results may suggest that the neurons, positive for both dopamine and GRP in VTA, project to the BLA (***Figure 6C***), and the *Grp* gene knockout affects these projections and the dopamine function in the BLA and as a result induces susceptibility in SEFL in the *Grp^-/-^* mice.

**Figure 6.**
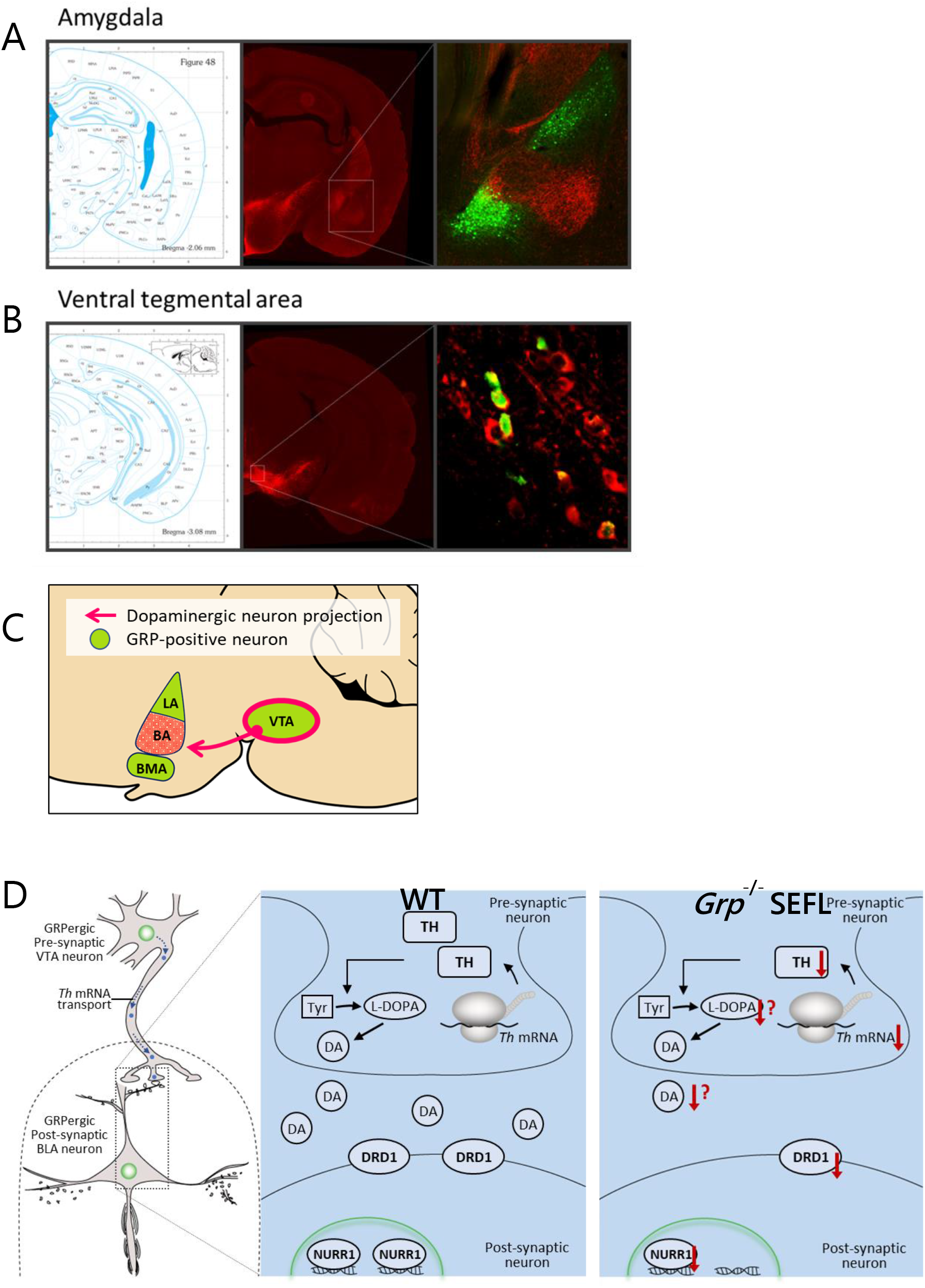
A possible relationship between the GRPergic circuit and dopaminergic circuit in extinction learning. The brains were harvested from 3-month old *Grp^-/-^* male mice and coronally sectioned at a thickness of 40 μm. Immunohistochemistry was performed with antibodies against the tyrosine hydroxylase (TH, red) and GFP (green). (A) Dopaminergic neurons project to the BA subregion of amygdala. GRP-positive cells in LA and BMA are not projected from the dopaminergic neurons. (B) The GRP is expressed within dopaminergic subpopulations of the VTA. (C) Schematic of the dopaminergic projections from the VTA to the GRP-positive regions. (D) A proposed model for the dopamine signaling in the GRPergic circuit in extinction learning after SEFL. The *Grp* gene knockout may affect the GRPergic/dopaminergic projections followed by decreased dopamine release and induce susceptibility in SEFL in the *Grp^-/-^* mice.

## Discussion

We generated the *Grp^-/-^* mice and showed that they are deficient in fear extinction in the SEFL behavioral paradigm, designed to model aspects of PTSD in humans. Retrograde tracing showed GRPergic connections between the BLA, mPFC and hippocampus, the areas critically involved in fear extinction. Transcription of several genes related to the dopamine signaling was downregulated in the BLA of the *Grp^-/-^* mice following the recall of fear memory in SEFL. These data point to the GRP and GRPergic cells as a molecular and neural-circuitry link to dopamine in regulating fear extinction. Moreover, the *Grp^-/-^* mice hold promise as an important genetic mouse model of PTSD-like symptoms.

The *Grp^-/-^* mice were generated with the cDNA encoding the green fluorescent protein (GFP) knocked in the first exon of the *Grp* gene locus. The presence of the GFP allowed for visualization of the learned fear circuitry by co-labeling the GRP-positive principal neurons and rAAV2-retro-based tracing (Tervo et al., 2016). The results demonstrate that the GRP-positive area in the BLA receives projections from the GRPergic cells of the TE3 area of the auditory cortex but not from the MGm/PIN area of the auditory thalamus. However, the TE3 receives the GRPergic projections from the MGm/PIN. Thus, the GRPergic circuitry is involved in processing of the auditory CS information via the multisynaptic indirect, but not the monosynaptic direct, pathway connecting the MGm/PIN and BLA (LeDoux, 2000; Pitkanen, 2000). Because the *Grp^-/-^* mice show extinction deficits in SEFL, it is plausible that the MGm/PIN->TE3->BLA pathway is preferentially involved in cued fear extinction compared to the MGm/PIN->BLA pathway. The GRP is present in other brain areas of the neural circuitry regulating fear extinction, in particular, in the ventral hippocampus, mPFC and VTA (Martel et al., 2012; Martel et al., 2016; Shumyatsky et al., 2002; Uchida et al., 2014). Thus, the GRP may serve as a functional biomarker of fear extinction.

The SEFL behavioral protocol includes a single session of acute mild stress (2 hours of restraint in a tube) followed by mild fear conditioning (one session of two CS-US pairings, 0.5 mA shock) seven days later, which in turn is followed by fear extinction four days later and extinction memory recall additional 24 days later (Sillivan et al., 2017). Therefore, the conditions of the training are mild and the testing extends across a relatively long time.

The *Grp^-/-^* mice exhibit increased susceptibility to PTSD-like behaviors: they have an enhancement in fear memory and deficiency in extinction in SEFL compared to unstressed but fear-conditioned *Grp^-/-^* mice and both stressed- and unstressed-fear-conditioned wildtype control mice.

An increase in post-shock freezing during fear conditioning is used to predict slower extinction and susceptibility to trauma in inbred mice in SEFL (Sillivan et al., 2017). Based on these criteria, we were able to separate both the *Grp^-/-^* and wildtype mice into stress-resilient and stress-susceptible groups.

Interestingly, the *Grp^-/-^* mice that did not undergo acute restraint stress still showed impairment in extinction learning, although only on day one of extinction. A single restraint stress session impaired extinction retrieval, but not fear conditioning in our experiments in wildtype mice as was reported in rats (Briggs & McMullen, 2020). This also confirms extensive literature showing that various types of both acute and chronic stress lead to extinction deficits (Maren & Holmes, 2016).

Naïve *Grp^-/-^* mice displayed normal anxiety and pain sensitivity, consistent with normal transcription of immediate-early genes (IEG) *c-Fos* and *Arc* in the amygdala of naïve *Grp^-/-^* mice. These behavioral and IEG data suggest that deficient extinction observed in SEFL in *Grp^-/-^* mice is likely to reflect changes in learning and memory processes rather than in anxiety in naïve state. This observation in the *Grp^-/-^* mice has implications for understanding what causes PTSD in humans. Work in humans suggests that impairments in fear extinction may cause PTSD or be a pre-existing condition (Maren & Holmes, 2016). Poor extinction was shown to be a consequence rather than a cause of PTSD in combat-exposed PTSD patients (Milad et al, 2008). However, impairments in extinction learning were shown to predict later risk for developing PTSD (Guthrie and Bryant, 2006). In addition to the importance of trauma exposure itself, work in rodents using various fear extinction protocols and transgenic mouse models supports the idea that genetic and neural-circuitry preexisting conditions predict and may cause deficits in fear extinction in humans.

Importantly, the deficiency in extinction in the *Grp^-/-^* mice is persistent as evident in the enhanced freezing during the recall test 24 days following extinction in SEFL. Confirming our finding that the GRP removal prolongs fear extinction in the *Grp^-/-^* mice, the GRP decreases fear memory reconsolidation when applied intraperitoneally immediately following recall in rats (Murkar, Kent, Cayer, James, & Merali, 2018). Also, GRP protein levels and *Grpr* mRNA levels are increased following treatment with drug docosahexaenoic acid, which leads to facilitation of fear extinction (Hashimoto, Hossain, Katakura, Mamun, & Shido, 2018). These findings support earlier observations that a removal of the GRP signaling *in vivo* leads to enhanced and prolonged fear memory and deficient fear extinction in *Grpr^-/Y^* mice (Chaperon et al., 2012; Martel et al., 2012; Shumyatsky et al., 2002). Also, a blockade of the GRPR using a GRPR antagonist in the rat dorsal hippocampus disrupts extinction of aversive memory in inhibitory avoidance (Luft et al., 2006). Therefore, our current data add to earlier research that demonstrates that a decrease in the GRP signaling leads to deficiency in fear extinction, whereas an increase in the GRP signaling improves fear extinction.

Previous work showed that both long-term changes in gene transcription and activity-dependent gene transcription are critical for fear conditioning and fear extinction (Ponomarev, Rau, Eger, Harris, & Fanselow, 2010; Sillivan et al., 2017; Uchida & Shumyatsky, 2018; Uchida et al., 2017). Transcription of IEG *c-Fos* and *Arc* was increased in the BLA of *Grp^-/-^* mice 30 min following training in single-tone fear conditioning compared to their wildtype counterparts, suggesting that the amygdala activity is abnormally enhanced following fear learning in *Grp^-/-^* mice. Similarly, there is a decrease of neuronal activity in the mPFC and an increase in the BLA following fear extinction in the *Grpr^-/Y^* mice (Martel et al., 2012).

To shed a light on the molecular mechanisms involved in recall of fear memory following extinction of fear, we examined transcription of several genes implicated in fear, fear extinction or PTSD in the BLA of the *Grp^-/-^* mice (Maren & Holmes, 2016; Sillivan et al., 2017; Uchida et al., 2017; Wingo et al., 2018). Transcription of several genes involved in the dopamine signaling was decreased following SEFL memory recall (two weeks after extinction). These genes encode for the tyrosine hydroxylase (TH), nuclear receptor related 1 protein (NURR1) and dopamine receptor D1 (DRD1). These results suggest that the GRP may be an upstream signaling molecule of the dopamine signaling involved in regulation of fear extinction.

Our data support previous work showing that dopamine is critically involved in regulating fear extinction. Recent work in humans shows that the presence of dopamine during consolidation of memory for fear extinction can lead to better outcomes during recall (Gerlicher et al., 2018). Enhancing dopaminergic signaling promotes rescue of deficient fear extinction (Bernardi & Spanagel, 2014; Whittle et al., 2016). Work in humans showed the role of the catechol-O-methyltransferase (COMT) enzyme activity, which degrades dopamine, in fear extinction (Lonsdorf et al., 2009; Norrholm et al., 2013; Panitz et al., 2018). Human carries of the 9-repeat (9R) allele of the striatum-enriched dopamine transporter 1 (DAT1), which may enhance phasic dopamine release, have improved extinction learning (Raczka et al., 2011). Recent work shows the importance of dopamine neurons in the VTA during fear extinction (Luo et al., 2018; Salinas-Hernandez et al., 2018). This work about the role of the VTA dopamine in fear extinction is interesting in light of our finding that the GRPergic cells are also dopaminergic in the VTA. Other work also showed that the *Grp* mRNA is a selective marker for dopaminergic subpopulations in the VTA and substantia nigra pars compacta (SNc) of the midbrain, both in mice and humans (Kramer, Risso, Kosillo, Ngai, & Bateup, 2018; Viereckel et al., 2016).

We found a decrease in the *tyrosine hydroxylase* (*Th*) mRNA levels, a critical enzyme involved in dopamine synthesis, in the BLA of the *Grp^-/-^* mice following SEFL recall. However, the TH protein is expressed in the VTA, and not in the BLA, based on our immunohistochemistry analysis. Thus, it is likely that the *Th* mRNA we isolated from the BLA, was located in synapses of the VTA presynaptic neurons projecting to the BLA. Indeed, *Th* mRNA is transported to axons in sympathetic neurons (Gervasi et al., 2016). Moreover, *Th* mRNA trafficking and local protein synthesis of the TH play an important role in the synthesis of dopamine in the presynaptic terminal (Aschrafi, Gioio, Dong, & Kaplan, 2017). It is also important to note that the changes in the *Th* mRNA levels are likely to be acute and result from the fear recall, as the TH protein levels were normal in naïve *Grp^-/-^* mice. Overall this finding supports the idea that the VTA projections to the amygdala are important for fear extinction, as the VTA is involved in both aversive and rewarding events and more specifically in acquisition and extinction of fear (Abraham et al., 2014; Luo et al., 2018; Salinas-Hernandez et al., 2018).

We found a decrease of the *dopamine receptor D1* (*Drd1*) mRNA, in the BLA of the *Grp^-/-^* mice following SEFL recall. In support of this finding, an activation of the substantia nigra dopamine neurons and the D1 receptors in the dorsal striatum during fear extinction prevents the renewal of fear (Bouchet et al., 2018). Also, a decrease in D1 expression was observed in the prefrontal cortex and amygdala of susceptible mice in chronic social defeat stress (Huang et al., 2016). Other work also show that dopamine receptor activation in the vmPFC, amygdala and NAc modulates fear extinction (Abraham et al., 2014; Hikind & Maroun, 2008; Holtzman-Assif et al., 2010; Mueller, Bravo-Rivera, & Quirk, 2010; Shi, Fan, Xue, Wen, & Zhao, 2017). Another paper showed the importance of the Drd2 neurons from the central amygdala using translating ribosome affinity purification (TRAP) technology in fear extinction (McCullough, Daskalakis, Gafford, Morrison, & Ressler, 2018). Prefrontal dopamine signaling via D1R and D2R is critical for extinction (Zbukvic, Park, Ganella, Lawrence, & Kim, 2017). Pharmacological activation of D1 receptors in the dorsal striatum did not impact fear extinction acquisition or memory, but blocked fear renewal in a novel context (Bouchet et al., 2018). These and other studies suggest that D1 and D2 receptor activation in the BLA is necessary for the acquisition and extinction of fear.

We also found a decrease in *Nurr1* mRNA levels in the BLA in our analysis. NURR1 is a transcription factor and an immediate-early gene located in the cell nucleus, thus, the changes we see in the NURR1 are in the BLA neurons. In Nurr1 KO mice, the normal development of midbrain dopamine neurons and the expression of dopaminergic phenotypic markers are disrupted (Castillo et al., 1998; Saucedo-Cardenas et al., 1998; Zetterstrom et al., 1997). NURR1 modulates dopamine signaling by increasing transcription of the *dopamine transporter* gene and the *tyrosine hydroxylase* gene (Sacchetti, Mitchell, Granneman, & Bannon, 2001; Sakurada, Ohshima-Sakurada, Palmer, & Gage, 1999). Since we found the *Th* mRNA levels decreased, it is possible that the decrease in the NURR1 led to the weakening of the *Th* gene transcription during fear memory recall. Thus, the GRP might be one of the upstream regulators of dopamine action in the extinction process. The role of the GRP in regulating dopamine function can be critical from the perspectives of dopamine involvement in both reward learning and fear extinction learning. This further suggests the GRP potential as a dopamine-oriented drug in psychotherapy approaches to maximizing consolidation of successful fear extinction and safety learning (Papalini, Beckers, & Vervliet, 2020).

Interestingly, our RNA-seq analysis provided additional support for the GRP KO mice to be a promising genetic model for PTSD as we found the *Ppm1f* gene to be significantly decreased in the GRP KO mice following SEFL in comparison to the wildtype mice following SEFL. The *Ppm1f* gene is regulated by stress in mice and is associated with anxiety, depression and PTSD in humans (Sullivan et al., 2019; Wingo et al., 2018).

Here, we have shown that the lack of the GRP leads to an enhancement in fear memory and deficient extinction of fear, when mild acute stress is combined with mild fear conditioning in SEFL behavioral paradigm. Earlier, using *in vivo* microdialysis, it was shown that acute restraint elicited the release of both corticotropin-releasing factor (CRF) and bombesin-like peptides (GRP is a bombesin-like peptide) in the central amygdala (Merali, McIntosh, Kent, Michaud, & Anisman, 1998).This and other accumulated evidence suggests that the GRP may be involved in integrating processing of stress and memory of fear (Roesler et al., 2014). Indeed, a recent paper, describing the *glucocorticoid receptor* gene knockout in the GRPergic neural circuitry, confirms our current findings regarding the importance of the GRPergic circuits in stress and fear memory integration (Inoue et al., 2018).

Overall, our work suggests that the GRP may be involved in fear extinction by regulating dopamine function. This work also suggests that the GRP-dopamine connection can be a promising molecular approach to decrease fear memory responses in therapy for fear- and anxiety-related disorders.

## Materials and methods

### Animals

*Grp^-/-^* mice were maintained on C57BL/6J background (N>10). The homozygous *Grp^-/-^* mice and their WT littermates were generated by breeding heterozygous *Grp^+/-^* mice, which in turn resulted from breeding of heterozygous mice to C57BL/6J mice (Jackson Laboratory). All mice were maintained on a 12-h light/dark cycle. Behavioral experiments were conducted during the light phase of the cycle, and mice were at least 12 weeks old at the time of training. This study was performed in strict accordance with the recommendations in the Guide for the Care and Use of Laboratory Animals of the National Institutes of Health. The Rutgers University Institutional Animal Care and Use Committee approved the protocol.

### Immunohistochemistry

Immunohistochemistry was performed as previously described (Martel et al., 2016; Uchida et al., 2014; Uchida et al., 2017). Mice were deeply anesthetized with Avertin (250 mg/kg i.p.) and transcardially perfused with 4% PFA. Their brains were postfixed overnight in 4%PFA and cryoprotected in 30% sucrose. The brains were sectioned (40μm) using a cryostat, and single or double immunofluorescence was performed on free-floating sections. Primary antibody is Rabbit polyclonal antibodies against green fluorescent protein (GFP) (1:500, Invitrogen). Secondary antibodies were conjugated with AlexaFluor-488 (1:500, Invitrogen). Images were acquired using an LSM 510 META laser confocal microscope or Observer Z1 (Zeiss) with multichannel excitation and detection options, including optimal factory-recommended filter configurations to minimize spectral bleed-through.

### rAAV2-based retrograde tracing

For virus injections, mice were anaesthetized intraperitoneally with avertin (250 mg/kg) and placed in a stereotaxic frame. The skull was exposed and a small portion of the skull over dorsal hippocampus was removed bilaterally with a drill. Subsequently, AAV vectors (rAAV2-CaMKII-tdTomato (Tervo et al., 2016); 1.0×10^13^ viral genomes per ml) dissolved in physiological saline were injected bilaterally into the LA (AP: −2.0 mm, ML: ±3.25 mm, DV: −4.25 mm) or BA(AP: −1.5 mm, ML: ±2.8 mm, DV: −4.75 mm) (0.25 ml volume). The needle was slowly withdrawn after 5 min. Mice were perfused 3 weeks after surgery and the brains were sectioned at a thickness of 40 μm. Anti-GFP immunofluorescence was performed on free-floating sections.

### Fear conditioning

Contextual and cued fear conditioning was performed as described previously (Shumyatsky et al., 2005). Mice were singly housed for at least 7 days prior the behavioral test. For CFC, each mouse was placed in the conditioning chamber (Med Associates) for 238 s before the onset of a 2-s foot-shock [0.75 mA, unconditioned stimulus (US)). After an additional 60 s in the chamber, the mouse was returned to its home cage. Twenty-four hours (LTM) or 1hour (STM) after training, the mouse was placed back in the chamber. Three hours later to test memory for cued fear conditioning, mice were placed in a novel environment in which the tone (120 s) that had been presented during training was given after a 1 min habituation period (pre-CS). The time spent freezing was assessed for 3 min using FreezeView software (Coulbourn Instruments).

### Pain sensitivity test

Response to the electric shock was assessed with naïve mice as described previously (Martel, Hevi, Kane-Goldsmith, & Shumyatsky, 2011; Shumyatsky et al., 2002). The intensity of the shock required for running, vocalization, and jump was determined for each mouse by delivering a 1-s-long shock every 30 s starting at 0.08 mA and increasing the shock 0.02 mA each time. Testing was stopped after all behaviors were noted.

### Open field test

This test was performed as previously reported (Martel, Hevi, Friebely, Baybutt, & Shumyatsky, 2010). The open-field consisted of a white arena (43.2cm×43.2cm×40cm) coupled to an automated video tracking system (Open Field Activity Software, Med Associates). Mice were placed in the corner of the arena, and the time spent in the peripheral area and the total distance traveled (locomotion) were measured. Results are expressed as the ratio of the time spent in the periphery over the total time spent in the arena.

### Elevated plus maze

This test was performed as previously reported (Martel et al., 2010). The elevated plus maze (1 m above the floor) consisted of a center platform (5cm×5cm), two open arms (40cm×5cm), and two closed arms (40 cm×5 cm) within walls (height 30 cm). Mice were placed individually in the center of the apparatus, and the time spent in each arm was measured for 10min using Limelight software (Coulbourn Instruments). Results are expressed as the percentage of the time spent in closed arms over the total time spent in the maze.

### Light-Dark transition test

A light/dark transition test was conducted as previously described. The apparatus used for this test comprised a cage (43.2cm×43.2cm×40cm) divided into two sections of equal size by a partition. One chamber was brightly illuminated, whereas the other chamber was dark. Mice were placed into the dark side of the cage at the start of the experiment and allowed to move freely between the two chambers for 10 minutes. The distance travelled in each chamber (cm) and time spent in each chamber (s) were recorded using automated video tracking system (Open Field Activity Software, Med Associates).

### Restraint stress

Restraint stress was performed as previously described by placing individual animals into clear 50 ml conical vials (Falcon Centrifuge Tubes) with ventilation holes for 2 hours. Tubes were placed flat in an open box in a biosafety cabinet with overhead lights on for the duration of the procedure. Mice in FC control groups were placed in a biosafety cabinet in another room during restraint stress and briefly handled in their home cages.

### Auditory fear conditioning and extinction

7 days after restraint stress, mice were exposed to training context A three times in one day for a total of 12 minutes to habituate them to the context. 24 hours later, mice underwent the following fear conditioning protocol: two minutes of exploration followed by two 30 second CS-US pairings that co-terminated with a 0.5 mA footshock (US) separated by a 60 or 120 second inter-tone interval (ITI; both produce the same results). The CS was an 85 dB, 10 kHz tone. This moderate protocol was used to avoid a ceiling effect in controls and the potential for induction of a depressive-like phenotype in stressed animals. Mice were removed from the training context 1 minute after the second shock and immediately returned to their home cages. Context A consisted of grid floors, a dim corner light in the room, no overhead lights, and 70% ethanol used for cleaning. Extinction training (4 days post shock) and remote memory retrieval tests (30 days post shock) were performed in novel context B, consisting of smooth plastic flooring, a plastic insert on the walls of the chamber, bright overhead lights, chamber lights on, orange scent, a 65 dB white noise and isopropanol for cleaning. Following a 2-minute exploration in Context B, animals were given 5 (recall) or 30 (extinction) CS presentations in the absence of the US (tone only), each separated by a 60 second ITI.

### RNA isolation and cDNA synthesis

Separate groups of mice were used to isolate RNA for qPCR and RNA-seq analyses. The amygdala was dissected as previously reported. In brief, mouse brains were immediately extracted and put on ice. Bilateral punches of the amygdala (preferentially including BLA) were obtained. Collected tissue was immediately put into RNA later (QIAGEN) until processing. Total RNA from dissected tissues was extracted by using the RNeasy Mini Kit (QIAGEN). For RNAseq, total RNA was sent to the company, which made the libraries, and the RNAseq were performed on illumina Novaseq6000 (Novogene). For qPCR, one microgram of total RNA was used for cDNA synthesis by SuperScript IV Reverse Transcriptase (Invitrogen). The cDNA was stored at −80°C until use.

### Quantitative real-time PCR

Real-time PCR was performed using the Applied Biosystems 7900HT Fast Real-Time PCR System with SYBR green PCR Master Mix (Applied Biosystems) according to the manufacturer’s protocol. PCR conditions were 10 min at 95°C, followed by 40 cycles at 95°C for 15 sec and 60°C for 30 sec. Amplification curves were visually inspected to set a suitable baseline range and threshold level. The relative quantification method was employed according to the manufacturer’s protocol in which all target mRNA expression levels were normalized to *Gapdh* expression.

### RNA-seq data analysis

Adapter removal and quality trimming of raw data was performed with fastp (Chen, Zhou, Chen, & Gu, 2018). Processed reads were then aligned with kallisto (Bray, Pimentel, Melsted, & Pachter, 2016) to the gencode vM25 mouse transcript sequences (Frankish et al., 2019). TPMs were reevaluated for each sample by first rounding the number of reads mapping to each transcript, then recalculating TPM. When gene level TPMs are presented, they are the sum of TPMs from each isoform of a gene. All analysis and graphs were produced with the R programming language (R Core Team, 2020) and the tidyverse set of packages (Wickham et al., 2019).

### Western blotting

Western blotting was performed (Uchida et al., 2017) using equal amounts of protein separated on 12% Bis-Tris gels (Life Technologies) and transblotted onto polyvinylidene difluoride membranes (GE Healthcare Bio-Sciences). After blocking with 5% skim milk, the membranes were incubated with anti-Tyrosine Hydroxylase antibody (MilliporeSigma) or anti-GFP Polyclonal antibody (Thermofisher). After incubation with HRP-linked anti-rabbit IgG, the blots were developed using the ECL-Plus Detection Kit (GE Healthcare Bio-Sciences). Densitometric analysis was performed using ImageQuant software (GE Healthcare) after scanning (Kwikquant imager).

### Statistics

Analyses of the data were performed using an appropriate ANOVA. Significant effects were determined using Fisher’s post hoc test or Bonferroni’s correction. Unpaired Student’s t tests were used for two-group comparisons. In all cases, p values were two-tailed, and the comparisons were considered to be statistically significant when p < 0.05. All data are presented as the mean ± SEM.

## Acknowledgments

This work was funded by grants from the National Institute of Mental Health (Grants No. MH107555 and MH080328 to GPS), the Brain and Behavior Research Foundation-NARSAD Independent Investigator Award, Whitehall, March of Dimes, The NJ Governor’s Council for Medical Research and Treatment of Autism, NJ Commission on Brian Injury Research and Busch (Rutgers) (to GPS). P.S. is supported by grants NIH (R35 GM124976), NSF (DBI 1936046) and subcontracts from NIH (R01 DK056645, R01 DK109714, and R01 DK124369), as well as start-up funds from the Human Genetics Institute of New Jersey at Rutgers University. IF is supported by PhD fellowship from CONACYT (Mexico). We thank XiaoXi Zhuang and Rene Hen for providing the pnCreFNF14.19 plasmid for gene targeting and Monica Mendelson and Zaiqi Wu for help with generating chimeric mice. We thank Mariana Salazar for the art work.

## Ethics

Animal experimentation: All experiments were approved and performed in accordance with Rutgers University Institutional Animal Care and Use Committee (IACUC) guidelines.

---

**Figure 1 – figure supplement 1.**
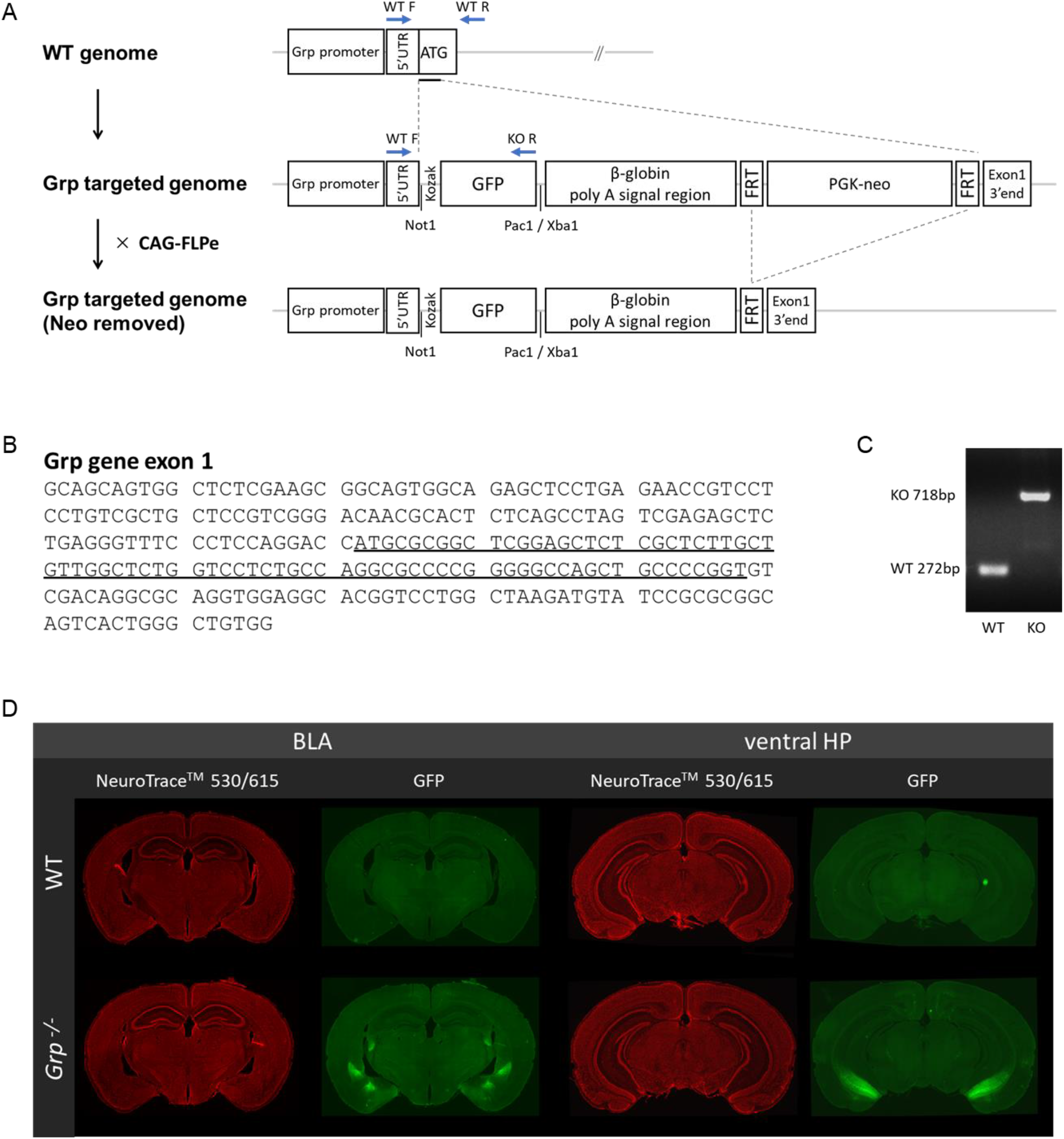
Generation of the *Grp^-/-^* mouse. (A) Diagram illustrating the *Grp* gene targeting design. PGK-neomycin was removed by crossing with CAG-FLPe mouse. The arrows show primers for PCR genotyping. (B) The underlined part show the DNA sequence deleted by targeting. (C) The following primers were used for genotyping PCR (WT F: GG ACAACGCACTCTCAGCCTAGT, WT R: AGACGGGGCTCCCTCTAGCTAG, KO R: ACTGGGTGCTCAGGTAGTGGTTGT). (D) The *Grp^-/-^* mouse shows no obvious anatomical abnormalities. Histology of the basolateral amygdala and ventral hippocampus in adult (3 months old) wildtype and *Grp^-/-^* mice. Consecutive 40-μm coronal sections were collected and stained for NeuroTraceTM 530/650 (1:100).

**Figure 2 – figure supplement 1.**
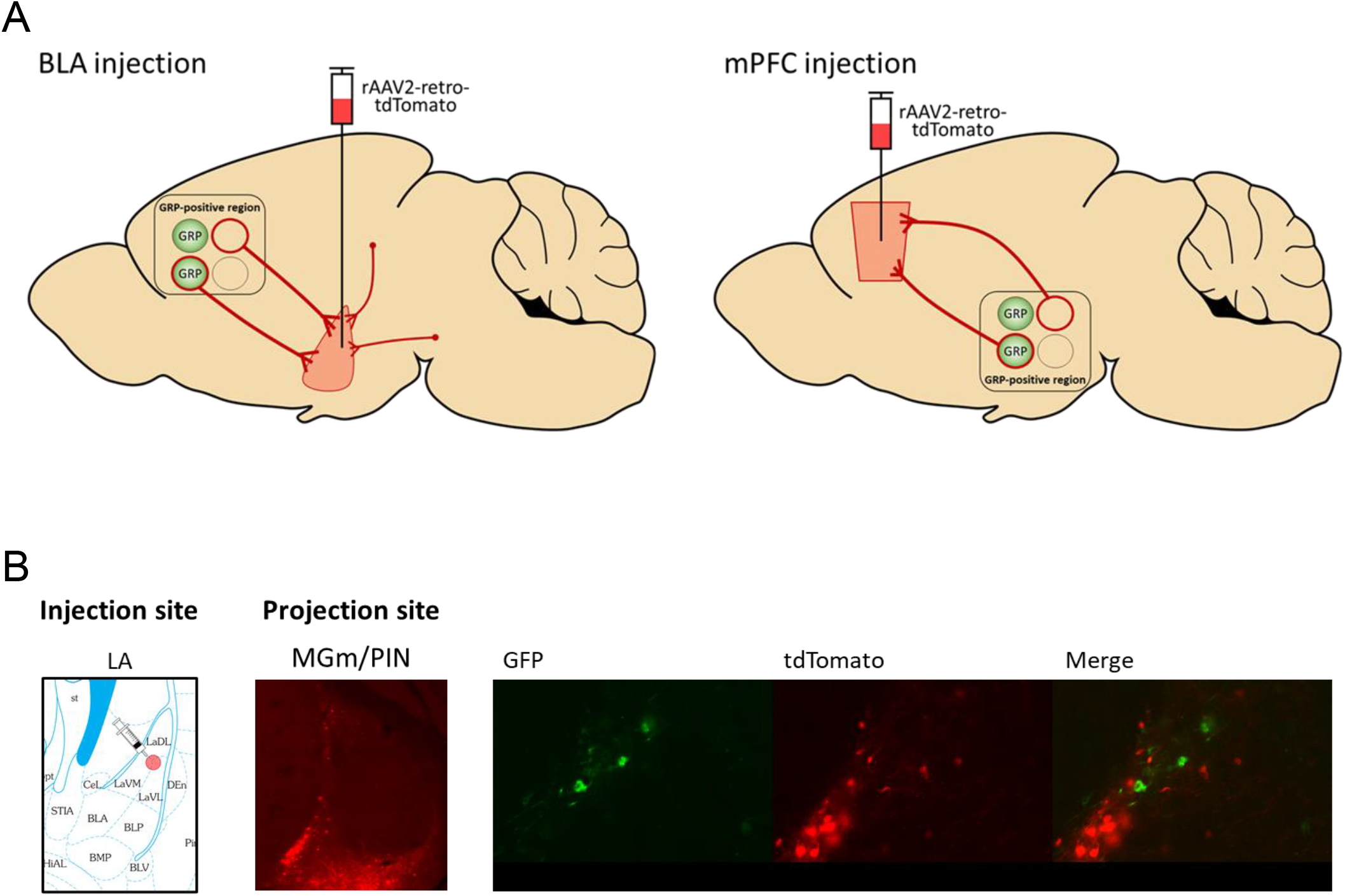
Schematic of retrograde tracing by AAV injection. Retrograde rAAV2-retro-CaMKII-tdTomato was injected into the BLA or mPFC of *Grp^-/-^* mouse brain. Three weeks following injections, the mice were perfused, and the brains were coronally sectioned at a thickness of 40 μm. (A) Schematic diagram of the virus injection. (B) rAAV2-retro-CaMKII-tdTomato was injected into LA. There were tdTomato-positive cells in the MGm/PIN, but there were no cells colocalized with GFP.

**Figure 3 – figure supplement 1.**
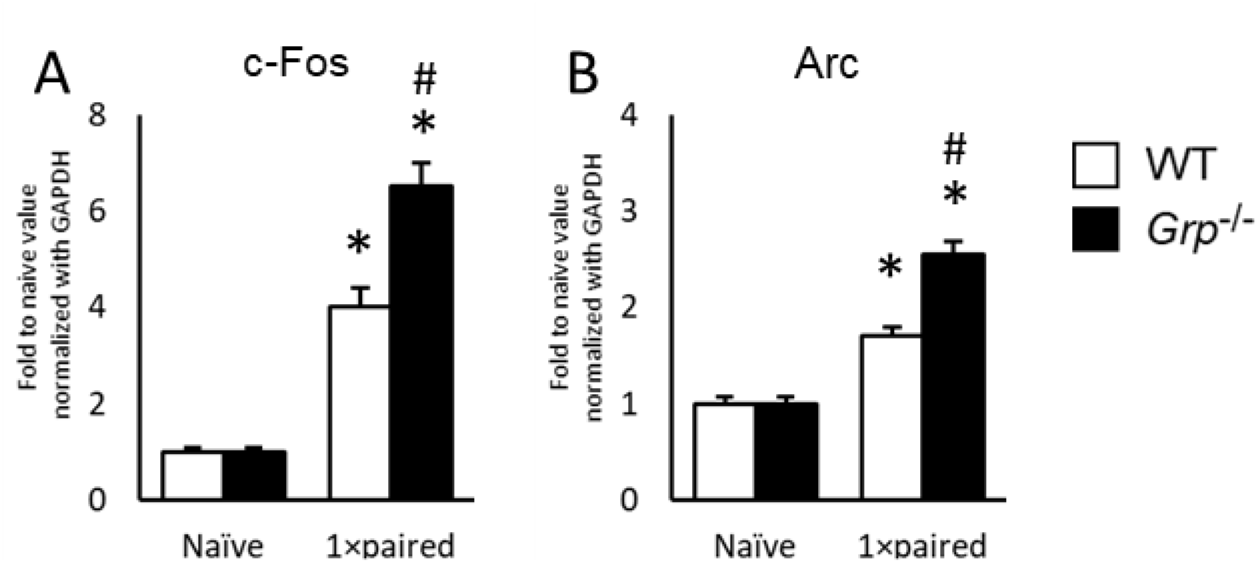
Transcription of immediate-early genes *c-Fos* and *Arc* in the amygdala following fear conditioning was enhanced in *Grp^-/-^* mice. (A) Expression levels of c-Fos mRNA in the amygdala of *Grp^-/-^* mice. (B) Expression levels of Arc mRNA in the amygdala of *Grp^-/-^* mice. The amygdala tissue was dissected 30 min after fear conditioning. c-Fos and Arc mRNA expression levels were normalized to Gapdh expression and verified by normalization to β-actin. Results are expressed as x-fold change normalized to WT naïve controls. *p<0.05 vs. respective naive. #p<0.05 vs. compared to wild type. Data presented as mean ±SEM.

**Figure 3 – figure supplement 2.**
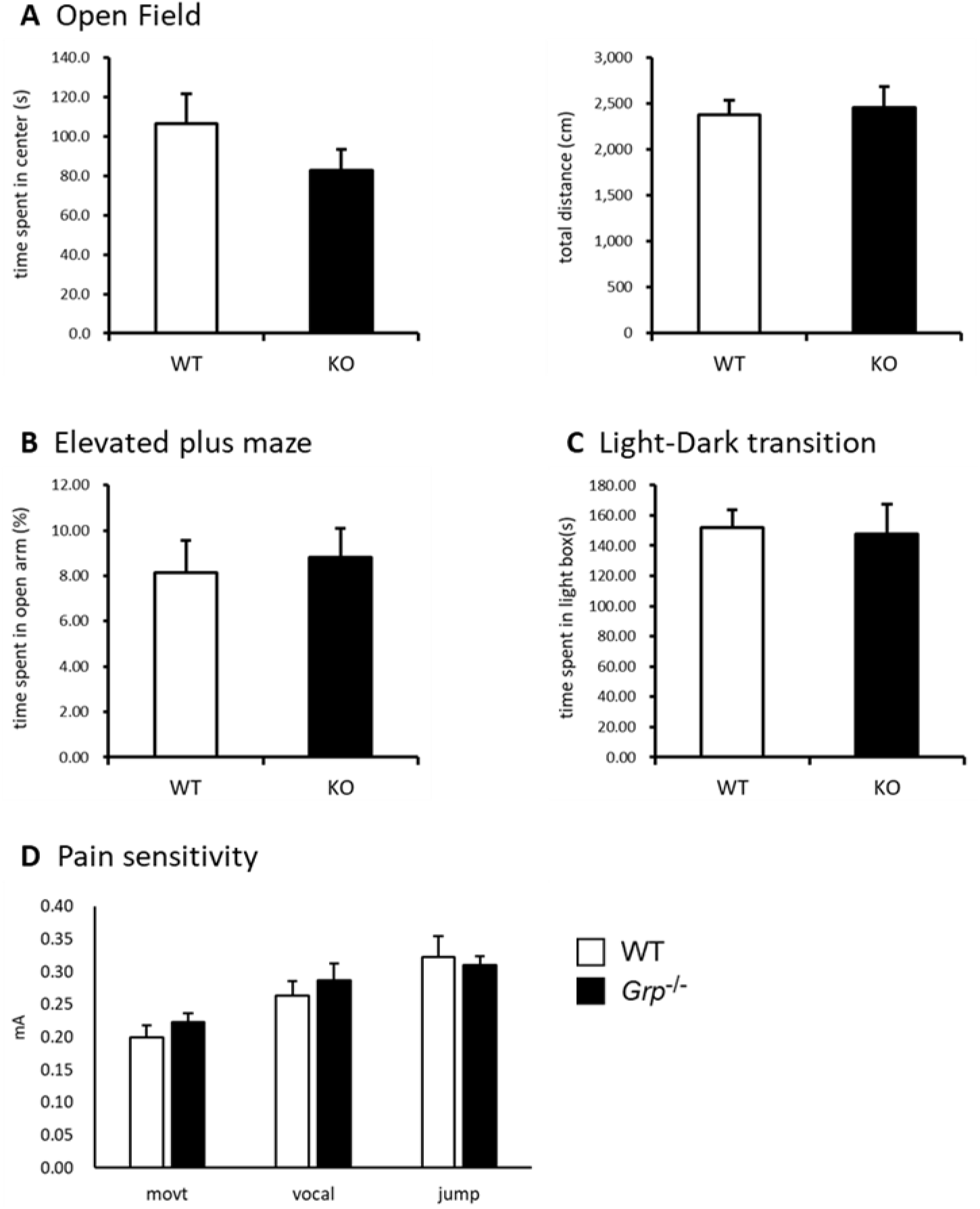
*Grp^-/-^* mice showed normal anxiety and pain sensitivity. (A) Open field test, (B) Elevated plus maze, (C) Light-dark transition test. No difference was found between groups (wildtype mice, n=15; knockout, n=15). (D) Pain sensitivity thresholds. The intensity of shock required to elicit three reactions, movement (movt), vocalization (vocal), and jump, were assessed and data are presented as the mean ±SEM. No difference was found between groups (wildtype, n=6; knockout, n=6).

**Figure 5 – figure supplement 1.**
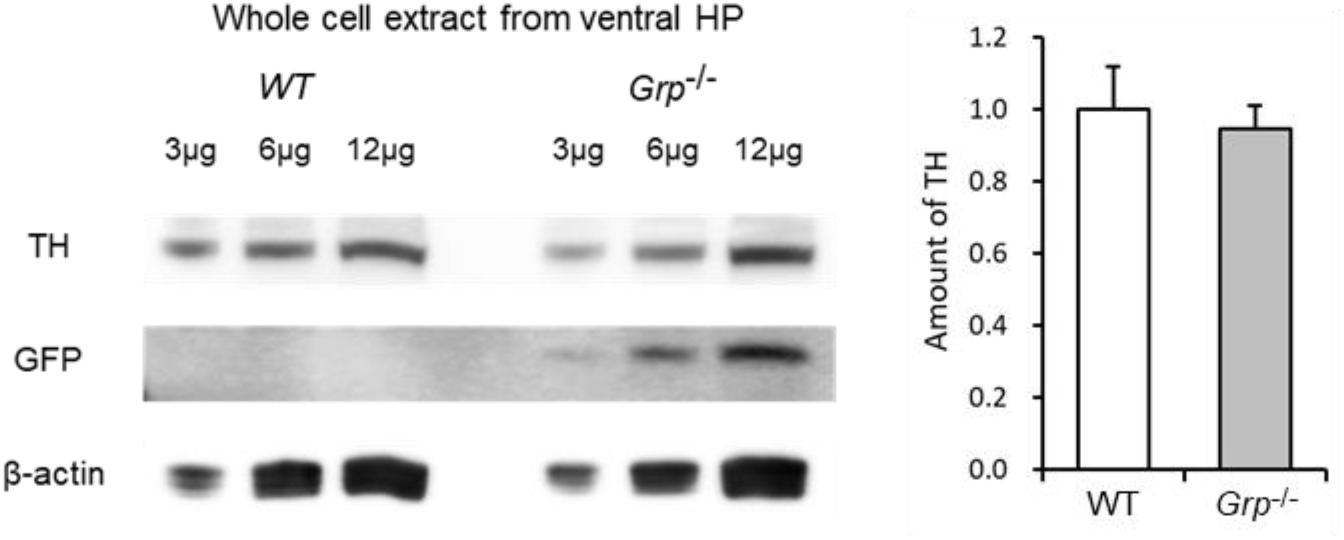
Tyrosine hydroxylase (TH) expression is normal in naïve *Grp^-/-^* mice. Western blot of whole-cell extracts from the ventral hippocampus of wildtype and *Grp^-/-^* mice, using antibodies against TH or GFP (the GFP cDNA is knocked-in into the *Grp* gene in the *Grp^-/-^* mice). 3, 6 and 12 μg of protein extract were separated on 12% Bis-Tris gels, and transblotted onto PVDF membranes. The relative expression level of the TH was normalized to β-actin levels.

**Figure 5 – figure supplement 2.**
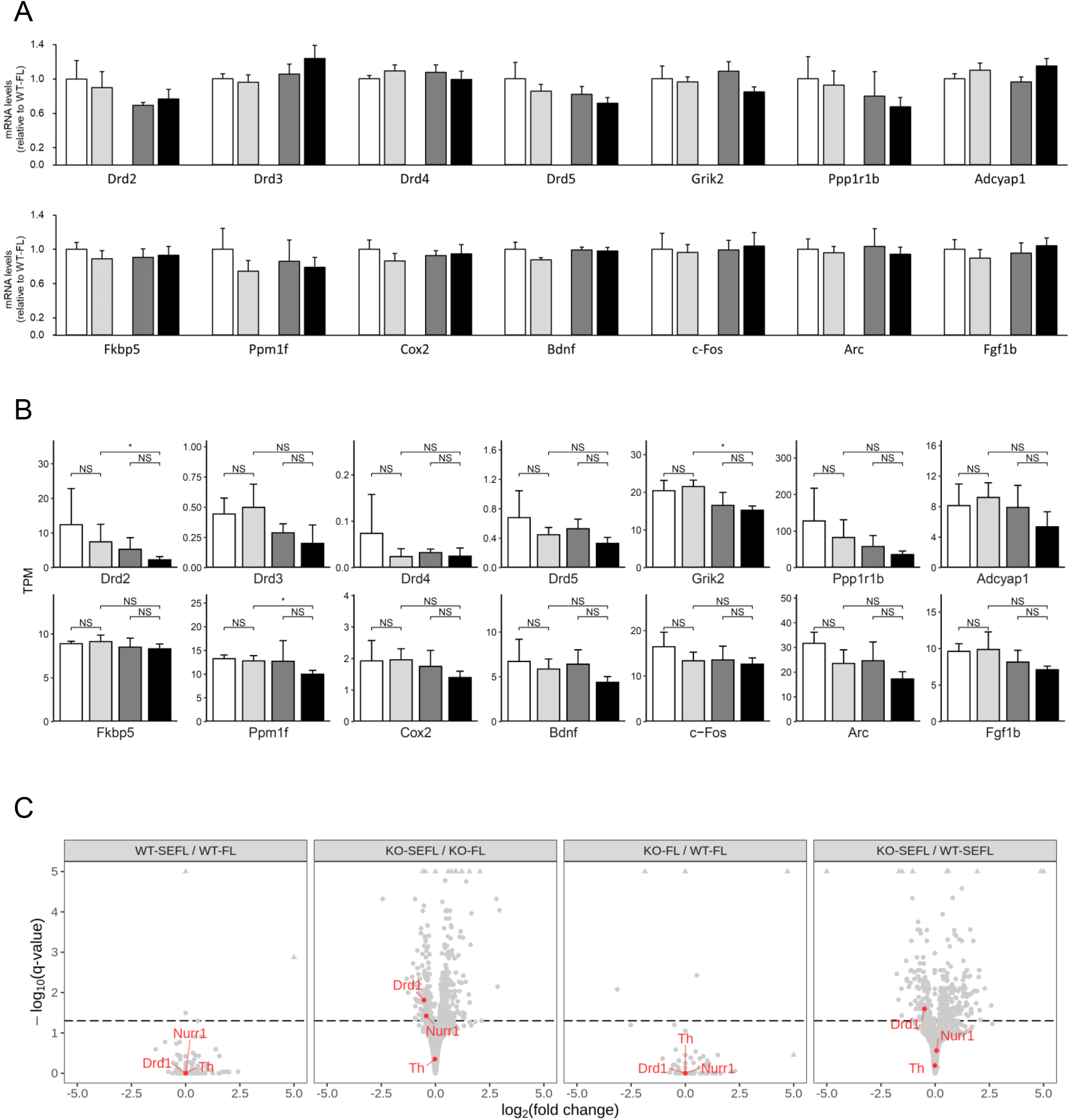
Dopamine signaling-related genes are downregulated in *Grp^-/-^* mice following recall in SEFL. (A-C) Analysis of dopamine signaling-related genes and stress susceptibility-related genes in the BLA. After the recall test, the mice were returned to their home cage and the amygdala tissue was dissected 30 min later. (A) Quantitative real-time PCR analysis. All target mRNA expression levels were normalized to *Gapdh* expression and verified by normalization to *β-actin*. Results are expressed as x-fold change normalized to WT controls. All measurements were performed in triplicate (WT-FL n=5, WT-SEFL n=11, KO-FL n=7, KO-SEFL n=10). (B) Expression differences in RNA-seq data recapitulate patterns observed in qPCR datasets, but some genes (*Drd2, Grik2* and *Ppm1f*) show significant differences in RNA-seq but not in qPCR. Comparisons of TPM (transcripts per million) based on 4 replicates and (*) indicates significant differences based on p-values <0.05 based on a t-test. (C) Volcano plot of differentially expressed genes based on RNA-seq datasets. 3 dopamine signaling-related genes are highlighted in red. The triangles are points where the fold-change or q-value (Benjamini-Hochberg corrected p-values of a t-tests), or both, are larger than indicated and have been reduced to aid in visualization. Namely, if the fold change was >5 or <-5, it has been changed to those values, and if −log10(q)>5, it was reduced to 5.

**Figure 5 – supplementary table 1.** mRNA expression of several genes involved in the dopamine signaling was differentially expressed in KO-SEFL as shown by two-way ANOVA followed by post-hoc test. Total number of each group (WT-FL n=12, WT-SEFL n=18, KO-FL n=13, KO-SEFL n=18) consists of two batches (1st batch: WT-FL n=5, WT-SEFL n=11, KO-FL n=7, KO-SEFL n=10, 2nd batch: WT-FL n=7, WT-SEFL n=7, KO-FL n=6, KO-SEFL n=8).

| Two-way factorial ANOVA | F value | Pr(>F) |
| --- | --- | --- |
| Genotype | 17.539 | <0.001 * |
| Stress | 16.928 | <0.001 * |
| Batch | 0.954 | 0.333 |
| Genotype:Stress | 8.420 | 0.0053 * |
| Genotype:Batch | 0.032 | 0.860 |
| Stress:Batch | 2.914 | 0.094 |

Nurr1
| Two-way factorial ANOVA | F value | Pr(>F) |
| --- | --- | --- |
| Genotype | 0.457 | 0.502 |
| Stress | 5.437 | 0.0235 * |
| Batch | 1.716 | 0.196 |
| Genotype:Stress | 0.515 | 0.476 |
| Genotype:Batch | 3.810 | 0.056 |
| Stress:Batch | 0.163 | 0.688 |

| Bonferroni's correction | P value |
| --- | --- |
| WT-FL vs WT-SEFL | 0.220 |
| KO-FL vs KO-SEFL | 0.041* |

Drd1
| Two-way factorial ANOVA | F value | Pr(>F) |
| --- | --- | --- |
| Genotype | 19.112 | <0.001 * |
| Stress | 0.010 | 0.921 |
| Batch | 1.600 | 0.211 |
| Genotype:Stress | 0.376 | 0.542 |
| Genotype:Batch | 0.005 | 0.944 |
| Stress:Batch | 0.734 | 0.396 |

| Bonferroni's correction | P value |
| --- | --- |
| WT-FL vs KO-FL | 0.0051 * |
| WT-SEFL vs KO-SEFL | <0.001 * |

